# Flexible analysis of transcriptome assemblies with Ballgown

**DOI:** 10.1101/003665

**Authors:** Alyssa C. Frazee, Geo Pertea, Andrew E. Jaffe, Ben Langmead, Steven L. Salzberg, Jeffrey T. Leek

## Introduction

A key advantage of RNA sequencing (RNA-seq) over hybridization-based technologies such as microarrays is that RNA-seq makes it possible to reconstruct complete gene structures, including multiple splice variants, from raw RNA-seq reads without relying on previously-established annotations [20, 32, 9]. But with this added flexibility, there are increased computational demands on upstream processing tasks such as alignment and assembly [28]. There is also a large and active community of developers contributing to downstream statistical modeling through the Bioconductor project [7]. However, there has been a gap between upstream processing tools and downstream statistical modeling tools that made it difficult to analyze assembled transcriptomes using Bioconductor. This gap has prevented rigorous statistical analysis of eQTL, timecourse, continuous covariate, or confounded experimental designs and has led to considerable controversy in the analysis of population-level RNA-seq data [4].

We have developed software that bridges the gap between transcriptome assembly and fast, flexible differential expression analysis (Supplementary Figure 1). First, our tool called *Tablemaker* uses a GTF file (output from any transcriptome assembler) and spliced read alignments to generate files that explicitly specify the structure of assembled transcripts, mappings from exons and splice junctions to transcripts, and several measures of feature expression, including FPKM (Fragments Per Kilobase of transcript per Million reads sequenced) and average per-base coverage (Supplementary Section 1). *Tablemaker* calls *Cufflinks* to estimate FPKM for each assembled transcript.

After the transcriptome assembly is processed with *Tablemaker*, it can be explored interactively in R using the *Ballgown* package. *Ballgown* reads *Tablemaker*’s assembly structure and expression estimates into an easy-to-access *R* object (Supplementary Section 2) for down-stream analysis. Alternatively, the *Tablemaker* step can be skipped, and the R object can be created from a transcriptome whose expression estimates have been calculated with RSEM’s `rsem-calculate-expression` [16]. Once the data has been loaded, *Ballgown* functions can be used to visualize the internals of a transcript assembly on a gene-by-gene basis, extract any of several relevant abundance estimates for exons, introns, transcripts, or genes, and perform straightforward linear-model-based differential expression analyses (Supplementary Section 3). The default statistical modeling framework included in *Ballgown* is flexible and computationally efficient, and its statistical significance and false discovery rate estimates exhibit better properties than *Cuffdiff2*’s (Figure 1a,b). These model comparisons are essentially equivalent to the modeling implemented in *limma* [27], though no empirical Bayes shrinkage has been implemented in *Ballgown*. However, almost any differential expression R package can easily be incorporated into the *Ballgown* workflow, since the main data structure provides easy access to feature-by-sample expression matrices, phenotype data, and paths to read alignment (BAM [17]) files.

**Figure 1.**
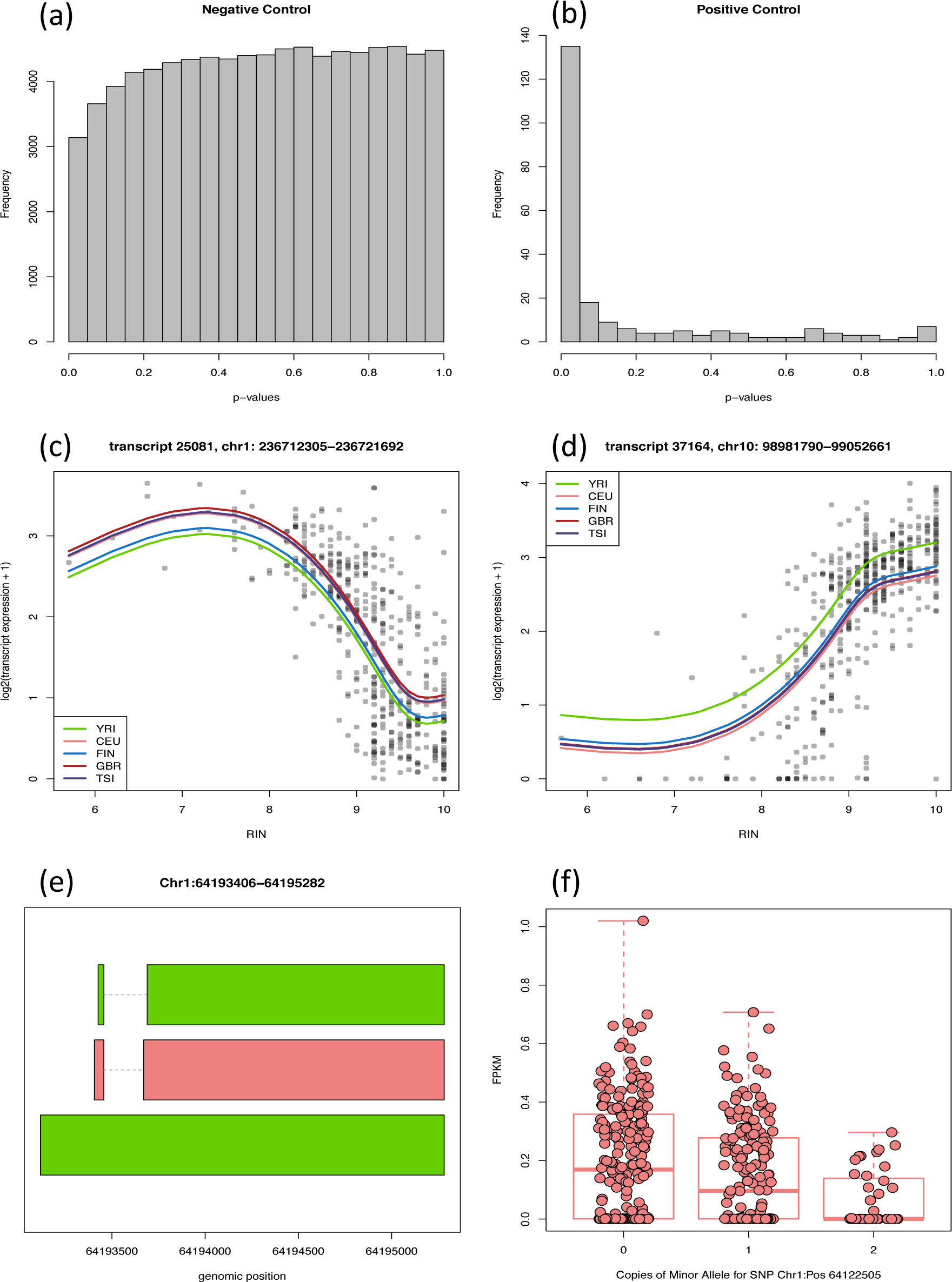
Experimental results obtained with the *Ballgown* framework. **a.** Distribution of transcript-level p-values obtained with *Ballgown*’s F-tests in an experiment without signal. **b.** Transcript-level p-value distribution from *Ballgown*’s F-tests for expression differences in Y-chromosome transcripts between males and females. **c-d.** Non-linear effects of RNA quality on transcript expression. These two transcripts (FDR *<* 0.001) and 1,497 others showed a relationship with RNA quality (RIN) that was significantly better captured by a non-linear trend with three degrees of freedom than a standard linear model. Colored lines shown are predicted values from a natural cubic spline fit and represent predictions for the specified population, assuming average library size. **e-f.** An assembled transcript that does not overlap any annotated transcripts but shows a significant eQTL. Panel **e** displays transcript structures for the locus in question; Panel **f** is a boxplot of the FPKM transcripts for the middle (red) transcript from panel **e**, which shows a consistent and statistically significant eQTL.

Here we illustrate the advantage of this approach using the widely used pipeline for transcript assembly, quantification, and differential expression analysis, the Tuxedo suite. This software suite aligns reads with *Bowtie* and *Tophat2* [11], assembles transcripts with *Cufflinks* [32] and performs differential expression analysis with *Cuffdiff2* [31]. This suite has been used in many influential projects [19, 34, 18], including the ENCODE [5] and modENCODE [10] consortium projects. However, *Cuffdiff2*’s statistical analysis capabilities are limited in terms of flexiblity, computational efficiency, accuracy of statistical significance and false discovery rate estimates. While several other fast, accurate tools for differential expression analysis like EdgeR [23], DESeq [2], and Voom [14] exist in the Bioconductor project [8], there is no existing infrastructure connecting these tools to assembly tools like the Tuxedo suite. Further, tools requiring per-feature read counts are often difficult to use for isoform-level analysis, since same-gene isoforms tend to have a high degree of overlap, and consequently, ambiguous read counts. We use *tablemaker* and *Ballgown* to integrate the Tuxedo suite with downstream statistical models in Bioconductor to improve statistical accuracy, flexibility of statistical modeling, and computational speed.

## Negative control experiment

First, we demonstrate that the default methods in *Ballgown* perform appropriately in a scenario where there is no differential expression signal. To create such a scenario, we downloaded and processed data from the GEUVADIS RNA sequencing project [13] [1] (Supplementary Section 4). After aligning RNA-seq reads, assembling the transcriptome, and processing the results with *Tablemaker*, we used *Ballgown* to load the data into R, where we extracted a single-population subset of data to study. The populations included in the GEUVADIS study were Utah residents with Northern and Western European ancestry (CEU), Yoruba in Ibadan, Nigeria (YRI), Toscani in Italy (TSI), British in England and Scotland (GBR), and Finnish in Finland (FIN). Considering only individuals in the FIN population (*n* = 95), we randomly assigned subjects to one of two groups and tested all assembled transcripts for differential expression between those two groups. We compared the results from using linear models (*Ballgown*), *Cuffdiff2*, and *EdgeR* [23] (at the exon level). For *Ballgown,* we used transcript FPKM as the transcript expression measurement, and we used per-exon read counts for EdgeR. In this type of experiment, the distribution of the *p*-values from all the transcripts should be approximately uniformly distributed, and *q*-values [29] should be quite high.

As expected, the transcript-level *p*-values from the linear model tests implemented in *Ballgown* were approximately uniformly distributed (Figure 1a), and all transcripts had *q*-values of approximately 1, indicating that these models are not overly liberal or prone to discovering signal where there is none. We compared this result to the statistical results from *Cuffdiff2* (version 2.2.1, the newest release available as of August 2014) on the same dataset, and we found that the *p*-values obtained using *Cuffdiff2* were not uniformly distributed, but that the distribution had more mass near 1 than near 0 (Supplementary Figure 5a). This indicates that *Cuffdiff2* may be somewhat conservatively biased, and calls into question the use of the *q*-value as a multiple testing adjustment, since it assumes uniformly-distributed *p*-values. Finally, at the exon level, *EdgeR* called two exons differentially expressed with *q* < 0.05, and the exon-level *p*-value distribution was not uniform, having a bit of extra mass around 0.1 (Supplementary Figure 5b). This result demonstrates that using a well-established, count-based methods gives a slightly too-liberal result on this kind of experiment, that *Cuffdiff2* is likely conservatively biased, and that using a linear model test like that implemented in *Ballgown* gives a reasonable *p*-value distribution without calling any transcripts differentially expressed.

For this experiment, the linear models from *Ballgown* took 18 seconds to run on a standard laptop (MacBook Pro, 8G memory). For comparison, *Cuffdiff2* took 69 hours and 148G of memory using 4 cores on a cluster node. *EdgeR* was also run on the laptop and took 2.5 minutes.

## Positive control experiment

The previous analysis showed that *Ballgown*’s default statistical tests are appropriately conservative when there is no signal present in the data. Here we present results from another idealized experiment to show that these default statistical tests are capable of making discoveries when differential expression really is present. For this experiment, we analyzed differential expression of Y-chromosome transcripts between males and females, so all transcripts should be differentially expressed. We again used the 95 FIN individuals in the GEUVADIS RNA-seq dataset (58 females, 37 males).

As expected, the *p*-value histogram from this experiment using the linear model framework implemented in *Ballgown* shows a very strong signal (Figure 1b). Of the 433 assembled transcripts on the Y chromosome, 225 had a mean FPKM greater than 0.01 in the males. Of those 225, 58% had *q*-values less than 0.05, and 72% had *q*-values less than 0.2, demonstrating that the models in *Ballgown* are capable of discovering true signal in the dataset. The *p*-value histogram for the latest *Cuffdiff2* version (2.2.1) does show some signal (Supplementary Figure 6), which is an improvement over earlier versions of *Cuffdiff2* on similar Y-chromosome tests [6]. However, only 29 of the 433 assembled transcripts were tested. Of those 29, 24 had *q* < 0.05 and 26 had *q* < 0.2, so *Cuffdiff2* 2.2.1 seems to perform adequately when tests are actually completed.

The Y chromosome linear models from *Ballgown* took less than 0.1 seconds to run after *Tablemaker*, and *Cuffdiff2* took 58 hours and 178G of memory on 4 cores. (Likely this footprint could have been substantially reduced by subsetting all BAM files and the merged assembly to only the Y chromosome, though this would require some extra processing time up front).

## Confirmation of statistical properties using inSilicoDB and simulated data

The experiments in the previous sections were designed to show that the default statistical modeling framework implemented in the *Ballgown* package is computationally efficient and gives reasonable results. However, those experiments do not necessarily represent realistic differential expression scenarios: usually some, but not all, features are truly differentially expressed between populations. We provide differential expression results from *Ballgown* and *Cuffdiff2* (versions 2.0.2 and 2.2.1) on two publicly-available experimental datasets (Supplementary Section 5.2). The first experiment [12] compared lung adenocarcinoma (*n* = 12) and normal control samples (*n* = 12) in nonsmoking female patients. The second experiment [33] compared cells at five developmental stages; we analyzed the data from two stages: embryonic stem cells (*n* = 34) and pre-implantation blastomeres (*n* = 78). On these datasets, the *p*-value distributions from the linear model tests implemented in *Ballgown* were reasonable, as were the *p*-value distributions from *Cuffdiff2* version 2.2.1, though *Cuffdiff2* 2.2.1 was more conservative than *Ballgown*. However, results from *Cuffdiff2* version 2.0.2 (downloaded from the InSilico DB database [3]) showed noticeable conservative bias (Supplementary Figure 7).

We also performed two simulation studies (Supplementary Section 5.3) to illustrate that a simple linear modeling approach like that implemented in *Ballgown* has better sensitivity and specificity than the *Cuffdiff2* approach.

## Analysis of RNA-seq experiments with complex designs

The *Ballgown* infrastructure also gives researchers the flexibility to explore the effects of using alternative expression measurements for analysis. There are two major classes of statistical methods for differential expression analysis of RNA-seq: those based on RPKMs or FPKMs, as exemplified by *Cufflinks*, and those based on counting the reads overlapping specific regions, as exemplifed by *DESeq* [2] and *edgeR* [23]. *Tablemaker* outputs both FPKM estimates from *Cufflinks* and average coverage of each exon, intron, and transcript (Supplementary Section 1). We investigated the effect of expression measurement using both simulated data and the GEUVADIS dataset, and we confirmed the expected result: differential expression results using average coverage and using FPKM were largely correlated (Supplementary Section 6). So in differential expression analyses, simple expression measurements like average per-base coverage may be sufficient.

An advantage of the *Ballgown* framework over *Cuffdiff2* is the added flexibility to compare any nested set of models for differential expression or to apply standard differential expression tools in *Bioconductor*, such as the *limma* package [26]. To demonstrate *Ballgown*’s flexiblity, we performed two popular analyses that have not been possible with standard transcriptome assembly and differential expression tools: modeling continuous covariates and eQTL analysis.

## Analysis of quantitative covariates

In the first analysis, we treated RNA Integrity Number (RIN) [24] as a continuous covariate [30] and used *Ballgown*’s modeling framework to discover transcripts in the GEUVADIS dataset [13] whose expression levels were significantly associated with RIN (Supplementary Section 4). Of 43,622 assembled transcripts with average FPKM above 0.1, 19,203 showed a significant effect (*q* < 0.05) of RIN on expression, using a natural cubic spline model for RIN and adjusting for population and library size [21].

A previous analysis of the GEUVADIS data modeled variation in RNA-quality as a linear effect [1]. We fit a model with a linear RIN effect and population and library size adjustments to each transcript, and we identified an enrichment of transcripts showing positive correlation between FPKM values and RNA-quality as expected (Supplementary Figure 3). To investigate the impact of using a more flexible statistical model to detect RIN artifacts, we tested whether a 3rd-order polynomial fit for RIN on transcript expression was significantly better than simply including a linear term for RIN after adjusting for population. We found that the cubic fit was significantly better than the linear fit (*q* < 0.05) for 1,499 transcripts (Figure 1c-d), suggesting that flexible non-linear models may be helpful when measuring the relationship between quantitative covariates and transcript abundance levels.

## Expression quantitative trait locus analysis

To demonstrate the flexibility of using the post-processed *Ballgown* data for differential expression, we next performed an eQTL analysis of the 464 non-duplicated GEUVADIS samples across all populations (Supplementary Section 4). We filtered to transcripts with an average FPKM across samples greater than 0.1 and removed SNPs with a minor allele frequency less than 5%, resulting in 7,072,917 SNPs and 44,140 transcripts. We constrained our analysis to search for *cis*-eQTLs where the genotype and transcript pairs were within 1000 kb of each other resulting in 218,360,149 SNP-transcript pairs. To adjust for potential confounding factors, we adjusted for the first three principal components of the genotype data [22] and the first three principal components of the observed transcript FPKM data [15]. The analysis was performed in 2 hours and 3 minutes on a standard Desktop computer using the MatrixEQTL package [25].

Visual inspection of the distribution of statistically significant results and corresponding QQ-plot indicated that our confounder adjustment was sufficient to remove major sources of bias (Supplementary Figure 4). We identified significant eQTL at the FDR 1% level for 17,276 transcripts overlapping 10,524 unique Ensembl-annotated genes. We calculated a global estimate of the number of null hypotheses and estimated that 5.8% of SNP-transcript pairs showed differential expression. 57% and 78% of the transcript-SNP called significant in the original analysis of the EUR and YRI populations [13], respectively, appeared in our list of significant transcript eQTL. 14% of eQTL pairs were identified for transcripts that did not overlap Ensembl annotated transcripts (Figure 1e-f).

## Computational Efficiency

The linear model differential expression testing framework built into *Ballgown* or *limma* provides computational benefits by taking advantage of the Bioconductor infrastructure. Supplementary Section 7 details timing and memory-use results for experiments presented here. These timing results show that *Cuffdiff2* is extremely computationally intensive, and since straightforward, flexible linear models are more accurate even for transcript-level analysis (Figure 1a-b, Supplementary Figures 5, 6, 8), a framework like *Ballgown* will drastically reduce the computational burden of differential expression analysis of assembled transcriptomes while still providing meaningful results.

## Summary

We have proposed the *Ballgown* suite as a bridge between upstream assembly tools and downstream statistical modeling tools in Bioconductor. The *Ballgown* suite includes functions for interactive exploration of the transcriptome assembly, visualization of transcript structures and feature-specific abundances for each locus, and post-hoc annotation of assembled features to annotated features. Direct availability of feature-by-sample expression tables makes it easy to apply alternative differential expression tests or to evaluate other statistical properties of the assembly, such as dispersion of expression values across replicates or genes. The *tablemaker* preprocessor writes the tables directly to disk and they can be loaded into *R* with a single function call. The *Ballgown*, *Tablemaker* and *Polyester* software packages are available from Bioconductor and GitHub (Supplementary Section 8), and code and data from the analyses presented here are available on GitHub (Supplementary Section 9).

## Acknowledgements

The authors would like to thank Peter A.C. ‘t Hoen and Tuuli Lappalainen for providing QC data and assistance with contacting ArrayExpress. JTL, GP, and BL were partially supported by 1R01GM105705 and AF is supported by a Hopkins Sommer Scholarship.

